# Plant diversity accurately predicts insect diversity in two tropical landscapes

**DOI:** 10.1101/040105

**Authors:** Kai Zhang, Siliang Lin, Yinqiu Ji, Chenxue Yang, Xiaoyang Wang, Chunyan Yang, Hesheng Wang, Haisheng Jiang, Rhett D. Harrison, Douglas W. Yu

**Author notes:** co-corresponding authors. DWY, 86 871 65110887; RDH, 86 871 65223377, Send proofs to DWY. Shared first-authorship (Kai Zhang, Siliang Lin).

## Abstract

Plant diversity surely determines arthropod diversity, but only moderate correlations between arthropod and plant species richness had been observed until Basset *et al.* (2012, Science 338: 1481-1484) finally undertook an unprecedentedly comprehensive sampling of a tropical forest and demonstrated that plant species richness could indeed accurately predict arthropod species richness. We now require a high-throughput pipeline to operationalize this result so that we can (1) test competing explanations for tropical arthropod megadiversity, (2) improve estimates of global eukaryotic species diversity, and (3) use plant and arthropod communities as efficient proxies for each other, thus improving the efficiency of conservation planning and of detecting forest degradation and recovery. We therefore applied metabarcoding to Malaise-trap samples across two tropical landscapes in China. We demonstrate that plant species richness can accurately predict arthropod (mostly insect) species richness and that plant and insect community compositions are highly correlated, even in landscapes that are large, heterogeneous, and anthropogenically modified. Finally, we review how metabarcoding makes feasible highly replicated tests of the major competing explanations for tropical megadiversity.

## Introduction

The relationship between plant diversity and insect diversity is fundamental to ecology because (1) it underpins global species estimates of arthropods based on plant diversity (Condon *et al.* 2008; Hamilton *et al.* 2013; Stork *et al.* 2015); (2) it improves our understanding of the drivers of arthropod diversity and assembly structure (Novotny *et al.* 2006; Lewinsohn & Roslin 2008; Pellissier *et al.* 2013); and (3) a strongly predictive relationship could open the way to using plant community metrics as surrogates for arthropod communities (and *vice versa*), thus improving the efficiency of efforts to conserve biodiversity, and ecosystem functions and services (Castagneyrol & Jactel 2012). In particular, arthropod species richness and community composition (i.e. species identities and frequencies) could serve as a sensitive method for detecting and quantifying the degree of forest degradation and recovery (Ji *et al.* 2013; Edwards *et al.* 2014), which is especially needed for the monitoring and verification of contracts to pay local populations and governments to protect and restore standing forest, also known as PES (Payments for Environmental Services) and REDD+ schemes (Reduction in Emissions from Deforestation and Degradation) (Bustamante *et al.* 2016).

*A priori*, plant diversity must surely predict insect diversity (Lewinsohn & Roslin 2008; Haddad *et al.* 2009), because insects depend directly (via herbivory, pollination, and housing) and indirectly (via consumption of herbivores) on plant species, and insect herbivores show dietary specialization to subsets of plant taxa (Novotny & Basset 2005). In addition, plant and insect coevolutionary interactions have driven the evolution of the vast diversity of plant and insect species today (Thompson 1994; Cruaud *et al.* 2012; Edger *et al.* 2015). (N.B. In practice, studies of terrestrial arthropod diversity tend to focus on insects, because insects make up the majority of described arthropods and can be easier to sample. This study will also follow this practice.)

Not surprisingly, many papers have reported significant correlations between arthropod (mostly insect) and plant beta and alpha diversities (reviews in Lewinsohn & Roslin 2008; Castagneyrol & Jactel 2012; Pellissier *et al.* 2013). In particular, work by Novotny *et al.* (2002, 2006, 2007) has strongly suggested that the primary driver of high species richness among herbivorous insects in tropical forests is simply the greater number of plant species in the tropics, rather than either higher levels of host specificity and beta diversity or more insect species per area of foliage. In short, the local number of insect species should increase nearly linearly with the local number of plant species, and with a slope greater than one, since each plant species is associated with multiple herbivore and predator species (Castagneyrol & Jactel 2012).

However, correlations between arthropod (mostly insect) and plant species-richness have shown only moderate fit. Castagneyrol and Jactel’s (2012) comprehensive meta-analysis reported mean correlations of only 0.39 and 0.51 for studies in single habitats and across multiple habitats, respectively, and a regression slope < 1, even for studies that focused on herbivores and pollinators.

Four possible reasons for this apparent lack of explanatory power are (1) geographic variation in the ratio of herbivores to plants and of non-herbivores to herbivores, due to coevolutionary and ecological interactions amongst plants, herbivores and their predators (Hamilton *et al.* 2013); (2) variation across plant species in their geographic ranges, which is positively correlated with total insect richness (Condon *et al.* 2008); (3) correlations and linear regressions being inappropriate models; and (4) incomplete taxon sampling (Lewinsohn & Roslin 2008). The last explanation is straightforward to test at least in principle. For instance, although Pellissier *et al.* (2013) successfully demonstrated a correlation between phylogenetic beta diversities of plant and butterfly communities, they also found that plant phylogenetic alpha diversity did not explain butterfly phylogenetic alpha diversity. One reason was that some of the local plant taxa were not consumed by Lepidoptera and therefore contributed to variance in plant alpha diversity but not to explanatory power. Presumably, those plant species are consumed by other insect clades, and a taxonomically more complete sample might have uncovered a positive relationship between plant and insect alpha diversity.

Thus, in a groundbreaking study involving 102 investigators and 129 494 arthropod specimens collected in twelve 0.04-ha quadrats of tropical forest (0.48 ha total), Basset *et al.* (2012) demonstrated that local tree species richness could predict the local species richness of both herbivore and non-herbivore arthropod taxa exceptionally well. For each of their eighteen taxon datasets (corresponding to ordinal or sub-ordinal guilds), Basset *et al.* (2012) used tree-species data from the 0.48 ha of sampling effort to extrapolate total arthropod species richness for the entire 6000-ha reserve. Overall, they found that what they called their “plant models,” which were parameterized species-accumulation curves that predicted the accumulation of arthropod species from the accumulation of tree species, were consistently able to predict “to a precision of 1%” independently derived best estimates of total arthropod species richness for the entire 6000-ha reserve.

In summary, Basset *et al.* (2012) showed that, given comprehensive taxon sampling and a more sophisticated statistical approach than correlations, plant species richness could indeed accurately predict arthropod species richness. However, due to their unprecedentedly huge sampling and taxonomic effort, Basset *et al.*’s (2012) protocol is effectively unrepeatable (and itself was unavoidably limited to a tiny area [0.48 ha]), but it would be highly desirable to be able to repeat this protocol efficiently in large numbers and over large spatial scales, i.e. to ‘operationalize’ the approach. At larger spatial scales (i.e. within and across landscapes), additional determinants of community structure can start to contribute, such as variation in environmental conditions and variation in regional species pools (Castagneyrol & Jactel 2012). It is at these larger spatial scales that plant community data would be most valuable in management for acting as a surrogate for arthropod diversity (and *vice versa*).

Metabarcoding is emerging as a promising way of advancing biodiversity research (Taberlet *et al.* 2012; Cristescu 2014). In metabarcoding, bulk samples of eukaryotes or environmental DNA are extracted, amplified, and sequenced for one or more taxonomically informative genes (DNA ‘barcodes’) (Taberlet *et al.* 2012; Yu *et al.* 2012; Ji *et al.* 2013; Cristescu 2014). Most importantly, despite false negatives (species failing to be detected) and false positives (falsely present species) being found in metabarcoding, due to primer bias (Yu *et al.* 2012; Clarke *et al.* 2014; Deagle *et al.* 2014; Piñol *et al.* 2015) and other errors in the metabarcoding pipeline (sequence errors and chimeras from PCR, library prep, and/or pyrosequencing, and species lumping and splitting in OTU clustering and taxonomic assignment), species richness and composition estimates from metabarcoded arthropod samples have been shown to correlate well with estimates calculated from standard morphological identification, even when the focal taxa are different (Yu *et al.* 2012; Ji *et al.* 2013; Edwards *et al.* 2014).

We therefore used metabarcoding to scale up Basset *et al.*’s (2012) approach, and we asked if plant diversity can predict insect diversity at landscape scales. Specifically, 1) does plant species richness predict insect species richness (alpha diversity); 2) does plant community composition predict insect community composition (beta diversity); and 3) is the predictive power of the plant model consistent across insect orders and over different seasons?

We report here that plant models parameterized with metabarcoding data produce landscape-scale estimates of total insect species richness that are very close to independent non-parametric estimates of insect species richness, and we also find a high degree of correlation between insect and plant community compositions in two widely separated tropical landscapes. As a result, we conclude that, armed with high-throughput methods, it should indeed be possible to operationalize Basset *et al.*’s (2012) important result that plant diversity can accurately predict arthropod diversity.

One potential benefit is that modern remote-sensing technologies, which show increasing promise at efficient assessment of vegetation (Féret & Asner 2014; Baldeck *et al.* 2015), might now also make possible the efficient management of a large proportion of animal biodiversity. Another benefit, and perhaps the most important one, is that it should now be possible to conduct highly replicated tests of the major competing explanations for tropical megadiversity (Lewinsohn & Roslin 2008), and we explain this in detail in the Discussion.

## Materials and Methods

### Study sites

We conducted our study in two montane landscapes in tropical southern China (Fig. 1), which differ in the nature of environmental heterogeneity they encompass and provide contrasting case studies of the relationship between plant and insect diversity at landscape scales.

**Figure 1.** Inventory plots in Yinggeling (*n* = 21) and Mengsong (*n* = 28). Colors stand for different elevation categories (orange = < 600 m; blue = 600 - 800 m; purple = ≥ 800 m) in Yinggeling, and for different habitat types (red = mature forest; green = regenerating forest; yellow = open land) in Mengsong. The Mengsong area left of the dashed line is included in Bulong Nature Reserve.

#### Yinggeling

Nature Reserve is located in central Hainan province (UTM/WGS84: 49N 328731 E, 2102468 N), a land-bridge island, and is the largest nature reserve in Hainan with an area of > 500 km^2^. The elevation ranges from 180 m to 1812 m, and the annual mean temperature correspondingly ranges between 24°C to 20°C. Mean annual rainfall is 1800– 2700 mm. The principal vegetation types are tropical montane rainforest and tropical montane evergreen broadleaf forest (Lin *et al.* 2013). Over 64% of the vegetation in the reserve is in a near-pristine state, although many of the carnivores have been extirpated (Lau *et al.* 2010).

#### Mengsong

(UTM/WGS84: 47N 656355 E, 2377646 N) is a sub-catchment of the upper Mekong River, with an area of ~100 km^2^. The elevation ranges from 800 m to 2000 m. Mengsong has a subtropical climate influenced by the Indian monsoon. The annual mean temperature is 18°C (at 1600 m asl). Mean annual rainfall varies between 1600–1800 mm, 80% of which falls in May–October. Mengsong has a > 200-year history of occupation by indigenous farmers, who formerly practiced swidden agriculture (Xu *et al.* 2009). Hence, today the landscape is a mosaic of mature forest with a history of selective cutting, forest that has naturally regenerated from clearance, and currently open land, such as terrace tea fields and grasslands. The principal primary vegetation types are seasonal montane rain forest in valleys, which grades into tropical montane evergreen broadleaf forest on upper slopes and ridges (Zhu *et al.* 2005). Part of Mengsong was included in Bulong Nature Reserve established in 2009. As with Yinggeling, many of the larger vertebrates have been extirpated (Sreekar *et al.* 2015).

### Biodiversity sampling

#### Yinggeling

Twenty-nine 50×50 m plots were set up in Yinggeling in May 2009 (10 plots) and September 2011 (19 plots). The plot locations were selected from a satellite image to incorporate as much of the substantial topographic variation found within the nature reserve as logistically possible. However, plot locations were not strictly randomly chosen. Plots established in 2009 were clustered, so for our study, only one plot was selected randomly from each cluster to minimize pseudo-replication. In total, 21 plots in Yinggeling were included (Fig. 1). All trees ≥ 5 cm DBH (‘diameter at breast height,’ which is set at 1.3 m from the soil surface) in each plot were surveyed. Species name, DBH, height and crown width were recorded. Field identifications were conducted by local experts, and trees that were not identified to the species level were excluded from further analyses.

Insects were sampled in the wet season (September to November 2011) using a Malaise trap located at the center of each plot for an average of 16 days (range: 12 - 25) depending on the weather, which affects capture efficiency. Malaise traps are widely used in sampling insect diversity, and are designed to capture flying insects that escape upwards (mainly Diptera and Hymenoptera), but are generally not suitable for Coleoptera, which drop when they hit a barrier (review in Russo *et al.* 2011). The collecting bottles on the Malaise traps were filled with 99.9% ethanol. Upon collection, the contents of each bottle were sieved to remove ethanol and placed in a storage bottle with fresh 99.9% ethanol. Between samples, the sieve and other equipment were rinsed with water and ethanol-flamed to prevent DNA cross-contamination.

#### Mengsong

Twenty-eight 100×100 m plots were set up from April 2010 to May 2011, based on a stratified random sampling design described in Paudel *et al.* (2015a; b) (Fig. 1). Plots covered a gradient from heavily disturbed shrubland and grassland (*n* = 6), through regenerating forest (*n* = 12) to mature forest (*n* = 10). Each plot consisted of nine 10-m radius subplots arranged on a square grid with 50 m spacing (Beckschäfer *et al.* 2013). All trees, bamboos, and lianas with ≥ 10 cm DBH were recorded within a 10-m radius of the subplot center, and all trees, bamboos, and lianas with 2-10 cm DBH were recorded within a 5-m radius of the subplot center. Species name, DBH, distance and angle to the subplot center were recorded. All herbs, ferns, and woody seedlings with < 2 cm DBH were surveyed within 1-m radius of the subplot center using a Braun-Blanquet coverage estimator (total coverage for each species was estimated visually and recorded using cover-abundance scale within six cover classes). Vouchers of every species in each plot were collected, and field identifications were confirmed (or adjusted) based on comparison to herbarium material at the Xishuangbanna Tropical Botanical Garden (HITBC). The vouchers were later deposited at the Kunming Institute of Botany.

Insects were collected with Malaise traps in five subplots (four corners and the middle subplot) over six days at the end of the wet season (late September-December 2010, hereafter wet season) and at the end of the dry season (April-June 2011, hereafter dry season). The collection and laboratory processing protocol were same as for Yinggeling. Subplot samples were pooled within each plot for further analyses.

### DNA extraction, PCR amplification, pyrosequencing, and bioinformatic analysis

Samples were prepared by using one leg from all specimens equal to or larger than a large fly (~5 mm length) and whole bodies of everything smaller, added with 4 mL Qiagen ATL buffer (Hilden, Germany) (20 mg/mL proteinase *k* = 9:1) per 1 g of sample, homogenized with sterile 0.25-inch ceramic spheres in a FastPrep-24^®^ system (MP Biomedicals, Santa Ana, CA, USA) set on 5 m/s for 1 min at room temperature, and incubated overnight at 56 °C. The genomic DNA was extracted with the Qiagen DNeasy Blood & Tissue Kit from 10% of the lysed solution, with ≤ 900 μL per spin column, and quality-checked using the Nanodrop 2000 spectrophotometer (Thermo Fisher Scientific, Wilmington, DE, USA). DNA was PCR amplified for the standard mtCOI barcode region using the degenerate primers, *Fol-degen-for* 5′-TCNACNAAYCAYAARRAYATYGG-3′ and *Fol-degen-rev* 5′-TANACYTCNGGRTGNCCRAARAAYCA-3′ (Yu *et al.* 2012). The standard Roche A-adaptor and a unique 10 bp MID (Multiplex IDentifier) tag for each sample were attached to the forward primer. PCRs were performed in 20 μL reaction volumes containing 2 μL of 10 × buffer, 1.5 mM MgCl_2_, 0.2 mM dNTPs, 0.4 μM each primer, 0.6 U HotStart Taq DNA polymerase (TaKaRa Biosystems, Ohtsu, Japan), and ~60 ng of genomic DNA. We used a touchdown thermocycling profile of 95 °C for 2 min; 11 cycles of 95 °C for 15 s; 51 °C for 30 s; 72 °C for 3 min, decreasing the annealing temperature by 1 degree every cycle; then 17 cycles of 95 °C for 15 s, 41 °C for 30 s, 72 °C for 3 min and a final extension of 72 °C for 10 min. We used non-proofreading Taq and fewer, longer cycles to reduce chimera production (Lenz & Becker 2008; Yu *et al.* 2012). DNA from each sample was amplified in three independent reactions and pooled to reduce amplification stochasticity. A negative control was included for each sample during PCR runs to detect contamination. For pyrosequencing, PCR products were gel-purified by using a Qiagen QIAquick PCR purification kit, quantified using the Quant-iT PicoGreen dsDNA Assay kit (Invitrogen, Grand Island, New York, USA), pooled and A-amplicon-sequenced on a Roche GS FLX at the Kunming Institute of Zoology. The 21 Yinggeling samples were sequenced on two 1/8 regions (one 1/8 region shared with other samples). The 28 Mengsong samples were sequenced on one whole run (four 1/4 regions, late September-December 2010: wet season) and two 1/4 regions (April-June 2011: dry season), respectively.

We followed an experimentally validated bioinformatic pipeline (Yu *et al.* 2012; Ji *et al.* 2013) to denoise, deconvolute, and cluster the reads into 97%-similarity Operational Taxonomic Units (OTUs). *Quality control*: Header sequences and low-quality reads were removed from the raw output in the QIIME 1.5.0 environment (split_libraries.py: -l 100 -L 700 -H 9 -M 2 -b 10) (Caporaso *et al.* 2010b). We removed any sequences < 100 bp. *Denoising, deconvoluting and chimera removal*: PyNAST 1.1 (Caporaso *et al.* 2010a) was used to align reads against a high-quality, aligned data set of Arthropoda sequences (Yu *et al.* 2012), and sequences that failed to align at ≥ 60% similarity were removed. The remaining sequences were clustered at 99% similarity with USEARCH 5.2.236 (Edgar 2010), a consensus sequence was chosen for each cluster, and the UCHIME function was used to perform *de novo* chimera detection and removal. A clustering step is required for chimera detection because chimeric reads are expected to be rare and thus belong to small clusters only. The final denoising step used MACSE 0.8b2 (Ranwez *et al.* 2011), which aligns at the amino acid level to high-quality reference sequences and uses any stop codons in COI to infer frameshift mutations caused by homopolymers. *OTU-picking and Taxonomic assignment*: Sequences were clustered at 97% similarity using CROP 1.33 (Hao *et al.* 2011). Each cluster of sequences represents a set of COI reads that are more similar to each other than to any other cluster, and is called an Operational Taxonomic Unit (OTU), which should approximate or somewhat underestimate biological species. OTUs were assigned taxonomies using SAP 1.0.12 (Munch *et al.* 2008), keeping only taxonomic levels for which the posterior probability was ≥ 80%. OTUs containing only one read (which tend to be PCR or sequencing errors and are uncertain presences [Yu *et al*. 2012; Ficetola *et al.* 2015]) or assigned to non-Arthropoda taxa were discarded.

### Statistical analysis

Analyses were mostly performed using R 3.2.2 (R Core Team 2015) and packages *BAT* 1.3.1 (Cardoso *et al.* 2015) and *vegan* 2.3-0 (Oksanen *et al.* 2015). We converted metabarcoding read numbers to presence/absence data before analyses, because read numbers are unlikely to reflect biomass or abundance (Yu *et al.* 2012). We first analyzed all Insecta-assigned OTUs together and then separately analyzed each Insecta order with ≥ 50 OTUs, including Coleoptera, Diptera, Hemiptera, Hymenoptera, Lepidoptera, and Psocoptera (the last for Mengsong only). More than 90% of OTUs were identified to order rank. We did not conduct analysis at family level, since less than half of the OTUs were identified to this rank. We tentatively included Arachnida from Mengsong (*n* = 84 and 83 OTUs for wet and dry season, respectively) in our analyses of species richness, although they are a by-catch of Malaise traps and hence may have a more stochastic pattern.

### Species richness

To test whether plant species richness can predict insect species richness estimated from metabarcoded Malaise-trap samples (i.e. Insecta OTU richness), we first calculated Pearson’s correlations (*cor.test* function) to allow comparison with the wider literature (Castagneyrol & Jactel 2012). We then applied the “plant model” approach of Basset *et al.* (2012), as follows:

First, the mean number of arthropod OTUs and of plant species found with each additional vegetation sampling plot (i.e. rarefaction curves) (*specaccum* in *vegan*) were calculated. To control for the order in which plots are added, we randomly subsampled the data without replacement (method = “random”, permutations = 9999).

Second, we used CurveExpert 1.4 (Hyams 2009; default settings, except maximum iterations = 1000) to fit functions to the relationship between the mean number of arthropod OTUs and the mean number of plant species found with each additional sampling plot. Following Basset *et al.* (2012), we used AICc to choose the best function from three candidates: linear, power, and Weibull functions, and called the best function the “plant model.” For comparison, we also chose the best function from a broader selection of 25 candidates, including ones used in other studies (Dengler 2009). These alternative candidates included linear (including quadratic fit and 3^rd^ degree polynomial fit), exponential, power, growth, sigmoidal and rational functions (Hyams 2009). We selected the top three functions based on AICc. Since statistical models offering a good fit to the data do not necessarily result in a robust species richness estimates (Basset *et al.* 2012), we again fitted the top three functions to a random subset of data (20 out of 28 plots for Mengsong, 15 out of 21 plots for Yinggeling) to check for robustness. Then we predicted the arthropod OTU richness at 297 tree species (the number of species in all the Yinggeling survey plots) or at 807 vascular plant species (the number of species in all the Mengsong survey plots) with these newly parameterized models, and we compared these predicted richnesses against the observed arthropod OTU richnesses in our metabarcode datasets. The best function was the one with the smallest absolute difference (similar to the “lowest error of extrapolation” in Dengler 2009). The results from three and 25 candidate functions proved similar, and we thus focused on the results from the first approach, using three candidate functions.

Third, we extrapolated the best function (i.e. the “plant model”) to the total plant species richness in the landscape to generate the plant model’s prediction of total arthropod species richness. In Yinggeling, there are 603 tree species known from the total reserve (Zhang *et al.* 2013). In Mengsong, no information on total vascular plant species richness is available, so we used non-parametric estimators to extrapolate from the plot data to total vascular plant species richness ( *alpha.accum* in *BAT*).

Fourth, we used non-parametric estimators to independently estimate total arthropod OTU richness in the landscape directly from the arthropod dataset ( *alpha.accum*in *BAT*), and we compared this extrapolation with the prediction from the plant model (‘same-site prediction’). Specifically, we calculated the explained variance (R^2^) when fit to a y = x model, and also calculated correlations (*cor.test*, method = “pearson”) for insect orders.

Note that there exists no ‘true’ biodiversity dataset to test against. Basset *et al.*(2012) used both statistical (best-fitting function with lowest error of extrapolation) and biological arguments (relevant surveys in the world with large sampling efforts) to get their best estimates of arthropod diversity. As no comparable surveys with metabarcoding techniques are available, we necessarily used non-parametric estimators, choosing those (Jackknife1, Jackknife2 and Chao) that have been shown to perform better than other estimators (Walther & Moore 2005; Hortal *et al.* 2006). Non-parametric estimators use the species abundance/occurrence relationships (e.g. the number of species occurring in only one or two sites throughout the samples) to estimate the total number of species (Hortal *et al.* 2006). We further applied a correction factor (*P*) for these non-parametric estimators to improve performance under conditions of low sampling effort, which is usually the case in arthropod surveys (Lopez *et al.* 2012). In Mengsong, the above approach was firstly applied to the whole landscape, and then separately to forests (mature and regenerating forest) and open lands. We also included only tree data to build the plant model in the Mengsong forests.

Finally, to evaluate the generality of our plant models, we used Yinggeling’s plant model to try to predict Mengsong insect diversity, and used Mengsong’s plant model to try to predict Yinggeling insect diversity (‘cross-site prediction’). Yinggeling and Mengsong are good candidates for such a test, as they are in the same zoogeographic region (Holt 2013) but are far from each other (~1000 km). However, Yinggeling and Mengsong have different landscape histories, and their vegetation was sampled differently. To maximize comparability, we used only the Mengsong plots (*n* = 16) located within the forest of Bulong Nature Reserve (~60 km^2^) (Fig. 1) and only included trees ≥ 5 cm DBH in each plot.

### Community composition

To test whether plant species compositions can predict insect species compositions, we could use Mantel tests, Procrustes analysis, or co-correspondence analysis, with each approach offering advantages and drawbacks (review in Gioria *et al.* 2011). We elected to use Procrustes analysis, because it is generally more powerful than Mantel tests and is more widely used than co-correspondence analysis, facilitating comparison with other studies. Procrustes analysis superimposes one ordination on top of another by minimizing the sum of the squared distances between points from the first to the second ordination. The probability of the fit is calculated by comparing the observed sum of squared distances against those from a null distribution obtained by repeated Procrustes fitting of permuted data (Oksanen *et al.* 2015). We used a non-metric multidimensional scaling (NMDS) ordination (*metaMDS* in *vegan*, distance = “jaccard”) of community composition data as the input data matrices for the Procrustes analyses (*protest* in *vegan*, symmetric = TRUE). Because the Procrustes analysis requires an identical number of axes in both ordinations, we constrained the number of axes to two (k = 2) for Yinggeling and four (k = 4) for Mengsong across all analyses. Initial exploratory analyses found that two/four axes were optimal for most groups (low stress and consistent results). Stress values ranged from 0.08 - 0.24. We also used these approaches to compare variation in community compositions among insect orders and between the two seasons in Mengsong.

## Results

### Species richness

The 21 Yinggeling samples produced 40 261 sequence reads, and the 28 Mengsong samples produced 519 865 and 253 025 reads for wet season and dry season, respectively. After bioinformatic processing, we obtained 1 995 Insecta OTUs from Yinggeling, and we obtained 2 946 Insecta OTUs from Mengsong, including 2 073 in the wet season and 2 215 in the dry season samples. None of the PCR negative controls detected sample contamination.

All the best-fit ‘plant models’ were Weibull functions, except for Hymenoptera in the dry season in Mengsong, which was a power function. All the ‘plant models’ exhibited very close fits to the non-parametric estimates of total OTU richness for insects as a whole (Insecta) and for individual orders (Coleoptera, Diptera, Hemiptera, Hymenoptera, Lepidoptera and Psocoptera) in both Yinggeling (Figs. 2 and S1, all Pearson’s r > 0.98) and Mengsong (Figs. 2 and S1, all Pearson’s r > 0.99). Similar results were obtained when we analyzed forests and open land separately in Mengsong (Figs. S2 and S3), and similar results were obtained when we used only trees to build the plant models in Mengsong forests (Fig. S4). In contrast, and consistent with the results compiled by Castagneyrol & Jactel (2012), simple Pearson correlations between insect OTU richness and plant species richness at the survey plot level were low (Yinggeling: all r ≤ 0.14; Mengsong: all r ≤ 0.29 for both wet and dry seasons).

**Figure 2.** Same-site predictions. Scatterplot of plant-model estimates versus non-parametric estimates of total OTU (Operational Taxonomic Unit) richness, for Arachnida (for Mengsong only), Insecta and insect orders that contained ≥ 50 OTUs. To quantify the goodness-of-fit of these two estimates, explained variances (R^2^) for insect orders were calculated from a y = x model (dashed line). Circles, squares and triangles stand for P-corrected versions of the Jackknife1, Jackknife2 and Chao estimators, respectively. Different colors stand for different taxa; only taxa absent from Yinggeling are labeled in the Mengsong figures. The plant-model functions were Weibull for all taxa in Yinggeling and Mengsong, except for Hymenoptera (dry season) in Mengsong, which was a power function. Note breaks in the axes.

The cross-site plant-model predictions (Yinggeling plant model predicting Mengsong Insecta OTU richness and *vice versa*) lay within an error of 2X for all Insecta and for three orders (Coleoptera, Hemiptera, Hymenoptera), but not for Diptera and Lepidoptera (Fig. 3). Given the observed scatter, all correlations were, not surprisingly, very low (Mengsong’s wet-season plant model predicting Yinggeling: all Pearson’s r < 0.1; Yinggeling’s plant model predicting Mengsong’s wet season: all Pearson’s r ≤ 0.1. Similar results were obtained when we used Mengsong’s dry-season data, Fig. S5. In all analyses, we excluded the Insecta points to avoid double counting.)

**Figure 3.** Cross-site predictions. Scatterplot of plant-model estimates versus non-parametric estimates of total OTU richness. Symbols as in Figure 2. Plant models from one landscape were used to predict non-parametric estimates in the other landscape. Shown here are the Mengsong wet-season results. Mengsong dry-season results are similar and shown in Figure S5. The shaded area encompasses a two-fold difference between the two estimates (y = 0.5x to y = 2x). The plant-model functions were Weibull for all taxa in Yinggeling and Mengsong, except for Lepidoptera in Mengsong, which was a linear function.

### Community composition

Community compositions in Insecta and plants were highly correlated in both Yinggeling and Mengsong (Fig. 4, Table 1). Correlations were reduced somewhat but were still high when we considered insect orders separately, likely reflecting the smaller sample size at this taxonomic level (Figs. S6 and S7, Table 1). High correlations were maintained even when we limited our analyses to only forests in Mengsong (Table S1).

**Figure 4.** Procrustes superimposition plots between plant and Insecta communities (9999 permutations), with the input of non-metric multidimensional scaling (NMDS) ordinations calculated from binary Jaccard dissimilarities (k = 2 axes used in Yinggeling, k = 4 in Mengsong). All Procrustes correlation coefficients (r) are significantly different from zero at *p* < 0.001 (Table 1). Each pair of points represents a census plot; solid points indicate plant data, and hollow points insect data. Colors as in Figure 1.

**Table 1.** Procrustes correlations among plant, Insecta, and individual insect order communities in (a) Yinggeling and (b) Mengsong (9999 permutations), with the input NMDS (non-metric multidimensional scaling) ordinations calculated from binary Jaccard dissimilarities (k = 2 axes used in Yinggeling, k = 4 in Mengsong). Correlations with plants are bolded. In Mengsong, where insects were sampled in two seasons, wet vs. dry-season Procrustes correlations are presented on the diagonal and are underlined, and the proportions of wet-season Operational Taxonomic Units (OTUs) that were also collected in the dry season are reported below as percentages.

| (a) Yinggeling |  |  |  |  |  |  |
| --- | --- | --- | --- | --- | --- | --- |
|  | Coleoptera | Diptera | Hemiptera | Hymenoptera | Lepidoptera | **Trees ($n = 297$)** |
| Insecta ($n = 1995$) | | | | | | **0.76\*\*\*** |
| Coleoptera ($n = 239$) | | 0.54** | 0.47* | 0.63*** | 0.43* | **0.44\*** |
| Diptera ($n = 848$) | | | 0.64*** | 0.57** | 0.53** | **0.73\*\*\*** |
| Hemiptera ($n = 205$) | | | | 0.45* | 0.39 | **0.60\*\*** |
| Hymenoptera ($n = 163$) | | | | | 0.55** | **0.56\*** |
| Lepidoptera ($n = 263$) | | | | | | **0.65\*\*\*** |

(b) Mengsong
|  |  | Insecta | Coleoptera | Diptera | Hemiptera | Hymenoptera | Lepidoptera | Psocoptera | Vascular plants<br>(*n* = 807) |
| --- | --- | --- | --- | --- | --- | --- | --- | --- | --- |
| Insecta | wet (*n* = 2073) | <u>0.75</u>*** |  |  |  |  |  |  | **0.78** *** |
|  | dry (*n* = 2215) | 60.6% |  |  |  |  |  |  | **0.75** *** |
| Coleoptera | wet (*n* = 375) |  | <u>0.45</u>* | 0.60*** | 0.58*** | 0.55*** | 0.43* | 0.52** | **0.53** *** |
|  | dry (*n* = 603) |  | 38.0% | 0.75*** | 0.70*** | 0.63*** | 0.51*** | 0.60*** | **0.63** *** |
| Diptera | wet (*n* = 414) |  |  | <u>0.52</u>*** | 0.67*** | 0.71*** | 0.63*** | 0.52*** | **0.71** *** |
|  | dry (*n* = 413) |  |  | 70.9% | 0.78*** | 0.61*** | 0.52*** | 0.50** | **0.65** *** |
| Hemiptera | wet (*n* = 435) |  |  |  | <u>0.67</u>*** | 0.58*** | 0.44* | 0.50** | **0.70** *** |
|  | dry (*n* = 370) |  |  |  | 71.1% | 0.61*** | 0.44* | 0.57*** | **0.77** *** |
| Hymenoptera | wet (*n* = 409) |  |  |  |  | <u>0.68</u>*** | 0.50** | 0.57*** | **0.64** *** |
|  | dry (*n* = 360) |  |  |  |  | 70.0% | 0.51*** | 0.61*** | **0.72** *** |
| Lepidoptera | wet (*n* = 78) |  |  |  |  |  | <u>0.33</u> | 0.40 | **0.46** ** |
|  | dry (*n* = 95) |  |  |  |  |  | 61.1% | 0.47** | **0.38** |
| Psocoptera | wet (*n* = 71) |  |  |  |  |  |  | <u>0.42</u> | **0.55** *** |
|  | dry (*n* = 63) |  |  |  |  |  |  | 85.7% | **0.52** *** |

\* $p < 0.05$ , \*\* $p < 0.01$ , \*\*\* $p < 0.001$ , non-significant values shown in gray, after adjustment for multiple tests ( $p.adjust$ , method = “fdr”)

In Mengsong, 61% of Insecta OTUs recorded in the wet season were also recorded in the dry season. Interestingly, community compositions remained highly correlated between these two seasons for all Insecta and for individual insect orders, except Lepidoptera and Psocoptera (Table 1), showing that despite turnover across seasons, the different insect species compositions contain the same ‘ecological information,’ meaning that they consistently reveal the persistent compositional differences between the different vegetation plots, which themselves reflect differences in microhabitats, food sources and histories.

Finally, community compositions were highly correlated between most pairs of insect orders in both Yinggeling and Mengsong, with the exceptions of Lepidoptera and Hemiptera in Yinggeling and Lepidoptera and Psocoptera in the wet season of Mengsong (Table 1). This suggests that different insect orders also contain similar ecological information about habitat differences. Again, these results were upheld even when analyzing only forests in Mengsong (Table S1).

## Discussion

Our study has demonstrated (1) a close fit between estimates of total insect species richness that have been derived from plant models and from non-parametric estimators (alpha diversity, Figs. 2 and S1) and (2) a high degree of correlation between insect community composition and plant community composition (beta diversity, Fig. 4). We replicated our results in two landscapes (Yinggeling and Mengsong), in two seasons in one of the landscapes (wet and dry in Mengsong), and across multiple insect orders (Figs. S6 and S7). Furthermore, we have extended the plant-model approach from tropical America to tropical Asia, from a homogeneous forest of 60 km^2^ to two heterogeneous, anthropogenically modified landscapes (~100 - 500 km^2^), and from a labor-intensive dataset of morphologically identified specimens to an efficiently processed dataset of metabarcoded samples.

Our results thus strongly support Basset *et al.*’s (2012) finding that plant species richness can be an accurate predictor of insect species richness in tropical forest, and we show that plant and insect species compositions are highly correlated. Also, given that the Mengsong plant model was able to predict Arachnida species richness (Figs. 2 and S2), we find some support for the broader conclusion that plant diversity can be an accurate predictor of arthropod diversity. Of course, it will be necessary to carry out taxonomically more comprehensive sampling to be able to support the last conclusion strongly.

We also found that plant models from one landscape could predict the species richnesses of Coleoptera, Hemiptera, Hymenoptera, and all Insecta in another landscape to within an error of 2X (cross-site predictions, Figs. 3 and S5). One clear reason for the error in the cross-site predictions is that our sampling effort and design were different across sites, due to unavoidable logistical constraints. For instance, in Mengsong, each quadrat was sampled with five Malaise traps, but in Yinggeling, only one trap was used.

Regardless, it seems likely that different landscapes will have differently parameterized plant models, and we argue below that this provides an opportunity for testing theories of tropical megadiversity. Note also that for biodiversity management and conservation, precise estimates of insect species richness are not necessary. It is already useful to know that plant species diversity can indeed be used as a close surrogate for insect diversity and *vice versa*. We propose that metabarcoded insect samples should be tested as a sensitive and perhaps early indicator of forest degradation.

### Malaise traps and metabarcoding

When resources are limited, which they always are, a feasible way to carry out arthropod diversity surveys at large scales is to combine mass trapping (here, Malaise traps) with a high-throughput taxonomic method (here, metabarcoding). Naturally, this places limitations on the informational content of the resulting datasets. Any given trap type can collect only a portion of total arthropod biodiversity (Russo *et al.* 2011), and the downstream processes of DNA extraction, PCR amplification, high-throughput sequencing, and bioinformatic processing will result in false negatives (‘dropout species’) and false positives (‘artefactual species’ created by PCR-induced sequence chimeras, and PCR, sequencing, and clustering errors) (Bohmann *et al.* 2014).

PCR primers and software pipelines have been developed to minimize these errors (as used in Yu *et al.* 2012), but it is more important to understand how to interpret metabarcoding outputs judiciously. Multiple studies (Yu *et al.* 2012; Ji *et al.* 2013; Yang *et al.* 2014) have shown using both mock and real biodiversity samples that, despite false negatives and positives, metabarcoding datasets are nonetheless reliable for estimating *community-level* metrics of alpha and beta diversity. In other words, the degree to which arthropod samples (and thus locations) differ in species richness and composition can be quantified with metabarcoding, which is precisely the requirement of our study. We were thus able to recapitulate Basset *et al.* (2012) in finding that Weibull-function plant models accurately predict insect communities.

However, we cannot directly compare the parameter values of our plant models with Basset *et al.*’s (2012) models for two reasons. Most importantly, we used only Malaise traps, which are designed to capture flying insects that escape upwards, whereas many beetles and nonflying species are missed. Basset *et al.’*s (2012) collections were more comprehensive. Less importantly, our species concept is based on COI sequence similarity, which will differ somewhat (but not hugely) from morphological concepts in the Arthropoda (e.g. Schmidt *et al.* 2015). In any event, the use of DNA barcodes as a major input to species delimitation is now mainstream (Ratnasingham & Hebert 2013; Riedel *et al.* 2013; Tang *et al.* 2014), and barcodes are advantageous because they more efficiently reveal cryptic species (Condon *et al.* 2008).

### Explaining tropical herbivore megadiversity

Lewinsohn and Roslin (2008) partitioned the causes of tropical herbivore megadiversity into four components: (A) more host plant species in the tropics combined with some level of host specificity, (B) more arthropod species per tropical plant species, (C) higher host specificity of tropical herbivores, and (D) higher rates of species turnover (beta diversity) within the same host species in the tropics. Studies by Novotny *et al.* (2002, 2006, 2007) in Papua New Guinea, with temperate-zone contrasts in Central Europe, have supported component A (more host plant species) over the other three components, whereas a compilation of feeding experiments by Dyer *et al.* (2007) has supported component C: higher host specificity in tropical species. Two important observations made by Dyer *et al.* (2007) are that broad host range in the temperate-zone is more obvious when more host plant species are surveyed and that different host plant species in the tropics show higher levels of insect community differentiation than do different temperate-zone host plants, suggesting higher host specificity in tropical insects.

Given our results here and elsewhere that metabarcoding can deliver reliable metrics of arthropod communities (Yu *et al.* 2012; Ji *et al.* 2013; Edwards *et al.* 2014; Yang *et al.* 2014), we suggest that metabarcoding can now be used to carry out the large numbers of surveys needed to test the four competing (and perhaps coexisting) explanations of Lewinsohn and Roslin (2008). Components A (more tropical plant species) and B (more tropical arthropod species per plant species) can be differentiated by parameterizing plant models along a tropical to temperate gradient. A is self-evidently true, but if B is important then we should observe a steeper slope of the plant model in the tropics. Following Dyer *et al.* (2007), components C (tropical insects having narrower host ranges) and D (more rapid spatial turnover in tropical insects) can be tested and differentiated by the relative contributions to beta diversity of changing host plant species and spanning geographic distance, in tropical and temperate habitats. Although in many parts of the world, DNA-barcode databases are not yet sufficiently comprehensive to be able to identify most insects to species level, it should be possible to use a combination of sequence matching (Ratnasingham & Hebert 2007) and phylogenetic placement (Matsen *et al.* 2010; Berger *et al.* 2011) to identify most specimens to at least family level, allowing differentiation of herbivores from non-herbivores in the near future.

## Acknowledgments

We are very grateful to the three reviewers and to the editor for comments and suggestions. We thank Mei Long for field assistance, Zhaoli Ding for sequencing, Philip Beckschäfer for providing the Mengsong GIS layer, and Mengsong village, Bulong Nature Reserve and Hainan Yinggeling National Nature Reserve for field support. Funding for fieldwork in Mengsong was provided by GIZ/BMZ (Project Nos. 08.7860.3-001.00, 13.1432.7-001.00) on behalf of the Government of the Federal Republic of Germany. CYY and DWY was supported by the National Natural Science Foundation of China (31400470, 41661144002), the Ministry of Science and Technology of China (2012FY110800), the University of East Anglia and its GRACE computing cluster, and the State Key Laboratory of Genetic Resources and Evolution at the Kunming Institute of Zoology (GREKF13-13, GREKF14-13, and GREKF16-09).

## Data Accessibility

DNA sequences: Genbank’s Short Read Archive (Accession numbers: SRP065001 and SRP065147) and Dryad doi: http://dx.doi.org/10.5061/dryad.37b53.

Bioinformatic script, OTU representative sequences, and OTU table with assigned taxonomies: Dryad doi: http://dx.doi.org/10.5061/dryad.37b53.

All input data sets to R and R script: Dryad doi: http://dx.doi.org/10.5061/dryad.37b53.

## Author Contributions

KZ, DWY, RDH, and HJ designed the study. KZ, RDH, SL, CXY, CYY and HW collected data, led by RDH and SL. KZ, YJ, CXY, XW performed the molecular experiments. KZ and YJ performed the bioinformatic analyses. KZ performed the statistical analyses and wrote the first draft of the manuscript. DWY and RDH contributed substantially to revisions.

## Supporting information

**Appendix S1** R Markdown output.

**Figure S1** Same-site predictions. Scatterplot of plant-model (25 candidate functions) estimates versus non-parametric estimates of total OTU richness. Symbols as in Figure 2. Note breaks in the axes.

**Figure S2** Same-site predictions. Scatterplot of plant-model estimates versus non-parametric estimates of total OTU richness in the (a) forests and (b) open land in Mengsong. Symbols as in Figure 2. The plant-model functions for (a) forests were Weibull for all taxa except Hymenoptera (dry season), which was a power function. The plant-model functions for (b) open land were power for all taxa except Coleoptera (wet season), which was a linear function. Note breaks in the axes. **Figure S3** Same-site predictions. Scatterplot of plant-model (25 candidate functions) estimates versus non-parametric estimates of total OTU richness in the (a) forests and (b) open land in Mengsong. Symbols as in Figure 2. Note breaks in the axes.

**Figure S4** Same-site predictions. Scatterplot of plant-model estimates versus non-parametric estimates of total OTU richness in the forests in Mengsong. Only tree data were included to build the plant model. Symbols as in Figure 2. The plant-model functions were Weibull for all taxa except Hymenoptera (dry season), which was a linear function. Note breaks in the axes.

**Figure S5** Cross-site predictions. Scatterplot of plant-model estimates versus non-parametric estimates of total OTU richness. Symbols as in Figure 3. Shown here are the Mengsong dry-season results. The plant-model functions were Weibull for all taxa in both Yinggeling and Mengsong.

**Figure S6** Procrustes superimposition plots between plant and insect order communities in Yinggeling. All Procrustes correlation coefficients (r) are significantly different from zero at *p* < 0.05 (Table 1). Symbols as in Figure 4.

**Figure S7** Procrustes superimposition plots between plant and insect order communities in (a) wet and (b) dry seasons of Mengsong. All Procrustes correlation coefficients (r) are significantly different from zero at *p* < 0.01 for wet season and at *p* < 0.001 for dry season except for Lepidoptera (Table 1). Symbols as in Figure 4.

**Table S1** Procrustes correlations among plant, Insecta, and insect order communities in the forests of Mengsong (*n* = 22) (9999 permutations), with the input of non-metric multidimensional scaling (NMDS) ordinations calculated from binary Jaccard dissimilarities (k = 4). The percentage calculations, and the bold and underlined statistics as in Table 1.

